# Line drawings reveal the structure of internal visual models conveyed by cortical feedback

**DOI:** 10.1101/041186

**Authors:** Andrew T. Morgan, Lucy S. Petro, Lars Muckli

## Abstract

Human behaviour is dependent on the ability of neuronal circuits to predict the outside world. Neuronal circuits make these predictions based on internal models. Despite our extensive knowledge of the sensory features that drive cortical neurons, we have a limited grasp on the structure of the brain’s internal models. Substantial progress in neuroscience therefore depends on our ability to replicate the models that the brain creates internally. Here we record human fMRI data while presenting partially occluded visual scenes. Visual occlusion controls sensory input to subregions of visual cortex while internal models continue to influence activity in these regions. Since the observed activity is dependent on internal models, but not on sensory input, we have the opportunity to map the features of the brain’s internal models. Our results show that internal models in early visual cortex are both categorical and scene-specific. We further demonstrate that behavioural line drawings provide a good description of internal model structure. These findings extend our understanding of internal models by showing that line drawings, which have been effectively used by humans to convey information about the world for thousands of years, provide a window into our brains’ internal models of vision.

## Introduction

The visual system’s ability to utilise internal models is of crucial importance for human behaviour. This ability allows us to understand complex environments based on limited visual information, which increases our chances of survival. A Palaeolithic hunter, for example, had to recognise predatory threats even when such threats were partially occluded by trees. Similar internal models across individuals also provide shared references, facilitating communication and common goals, both of which are signatures of human behaviour.

Cortical neurons receive internal models and sensory signals as two separate sources of input. Neurons in early visual cortex are sensitive to sensory stimulation from only small portions of stimulus space, as defined by classical receptive fields. Cortical neurons also receive internal models through feedback (top-down) and lateral connections, which amplify and disamplify responses to the feedforward signals based on context^1,2^.

Although internal models are of crucial importance for human behaviour, they are challenging to study directly because their detection requires internal models to be disentangled from co-occurring sensory inputs. One strategy to access internal models during perception is to use visual occlusion to homogenise sensory input to a subsection of cortex. Functional brain imaging studies indicate that cortical feedback provides contextual filling in to patches of early visual cortex that process occluded scene information^3–8^. Here, we aim to determine the features that contribute to activity patterns associated with cortical feedback, thereby revealing the structure of the brain’s internal models involved in visual perception.

Category and depth are two candidate features of internal models that have been shown to modulate V1 responses^9,10^. These features can be quickly derived from scene statistics alone^11,12^, so both types of information could be present in V1 responses due to feedforward processing and need no feedback of internal models. However, to derive category and depth information from scenes requires scene information to be integrated over larger areas of the visual field than would be expected from direct communication amongst early visual cortical neurons. Therefore, we would expect internal models to play an integral role in the coding of category and depth in early visual cortex. In line with this hypothesis, previous work has suggested that the visual system rapidly determines abstract global features and then transmits them through feedback pathways to aid in early visual processing^13–15^. In this study, we investigated whether internal models delivered to early visual cortex carry category and depth information. To do so, we partially occluded visual scenes viewed by subjects during an fMRI experiment and attempted to read out category and depth information from responses in occluded portions of V1 and V2.

Additionally, we asked subjects for behavioural samples of their scene-specific internal models by way of sketched line drawings of occluded portions of visual scenes. Line drawings have remained largely unchanged from those made on the walls of caves by Palaeolithic hunters some 40,000 years ago. This consistency means that ancient cave drawings still effectively convey elements of the artist’s environment, even thousands of years later. In light of this, it has been suggested that line drawings embody fundamental components of how our visual systems represent the world^16,17^. If so, we expect behavioural drawings to depict internal models of missing scene information. We investigate this possibility by modelling occluded brain activity using line drawings.

Our data reveal that internal models sent to V1 and V2 contain category, but not depth information. Internal models also contain scene-specific orientation information that resembles behavioural predictions of missing visual scene information from line drawings. By combining functional brain imaging with a visual-occlusion paradigm and behavioural measures, we thus demonstrate that line drawings provide a window into the models that the brain creates internally.

## Results

We blocked feedforward input to subsections of retinotopic visual cortex during an fMRI experiment using a uniform visual occluder that covered one quarter of the visual field^4^ while participants viewed 24 real-world scenes. We localised subsections of V1 and V2 that responded either to the occluded portion of the visual field (lower-right image quadrant), or non-occluded visual field (upper-right and lower-left quadrants; Figure 1). This process yielded three regions of interest (ROIs) in each of V1 and V2, hereafter referred to as Occluded and Non-Occluded (either Upper-Right or Lower-Left), totalling six ROIs (3 positions in V1 and 3 positions in V2). We also mapped population receptive field (pRF) locations of individual voxels^18^ to ensure that their response profiles were within the ROIs in the occluded visual field.

**Figure 1.**
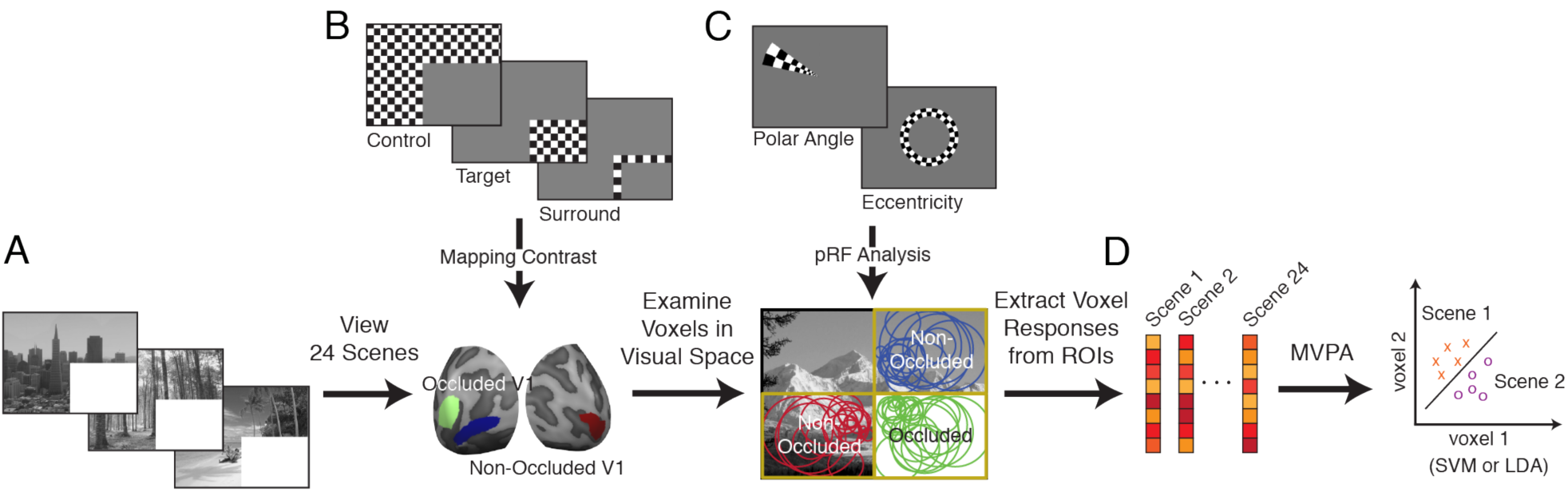
Experimental procedures. (A) Participants viewed 24 scenes with their lower right quadrants occluded. Scenes spanned 6 categories (Beaches, Buildings, Forests, Highways, Industry, and Mountains) and 2 depths (Near and Far). (B) Occluded and Non-Occluded subsections of early visual cortex were localised using mapping contrasts. (C) Retinotopic mapping data were used to separate V1 and V2 and to map population receptive fields (pRFs). Voxel pRFs not completely contained by the quadrant of interest (2σ from pRF centre) were excluded from further analyses. (D) Remaining voxels were included in multi-voxel pattern analyses (MVPA).

In this experimental design, Occluded V1 and V2 neurons receive homogenous white, non-diagnostic feedforward visual input (see Figure 1). Thus, any change in activity pattern is related to non-feedforward input to these areas. This input could be in the form of cortical feedback, where neighbouring neurons receive feedforward input, which is sent up the visual hierarchy and subsequently fed back to these Occluded portions of V1 and V2. Alternatively, Occluded neurons could receive information laterally through horizontal connections. The conservative mapping and large cortical area of Occluded ROIs in this study minimises this second possibility. Thus, activity patterns in Occluded V1 and V2 are related to internal models transmitted to the early visual cortex via cortical feedback with only a minimal contribution coming from lateral connections. To determine whether scene category and depth were related to internal models, we first attempted to decode these two scene characteristics from Occluded V1 and V2 responses. Scenes included in this study were therefore balanced across six categories (Beaches, Buildings, Forests, Highways, Industry and Mountains) and two spatial depths (Near and Far). To explore which scene-specific features are present in internal models, we also compared occluded responses to behavioural predictions of occluded scenes from drawings.

### Decoding high-level scene features from cortical feedback signals to early visual cortex

To measure the relationship of Category and Depth with V1 and V2 processing, we used single-trial, linear Support Vector Machine (SVM) classification, which has previously been shown to be sensitive in detecting cortical feedback^3,4^. Initially, we looked at the Non-Occluded V1 and V2 responses, which contain a mixture of feedforward input, cortical feedback, and lateral interactions. For Non-Occluded areas, we expected to find the strong decoding of all three types of information in our data, matching what has been found previously^3,4,9,10^. In our data, we indeed could decode individual scene, category and depth information in these areas (Figure 2a; one-sided Wilcoxon Sign-Rank, all p-values < 0.001).

**Figure 2.**
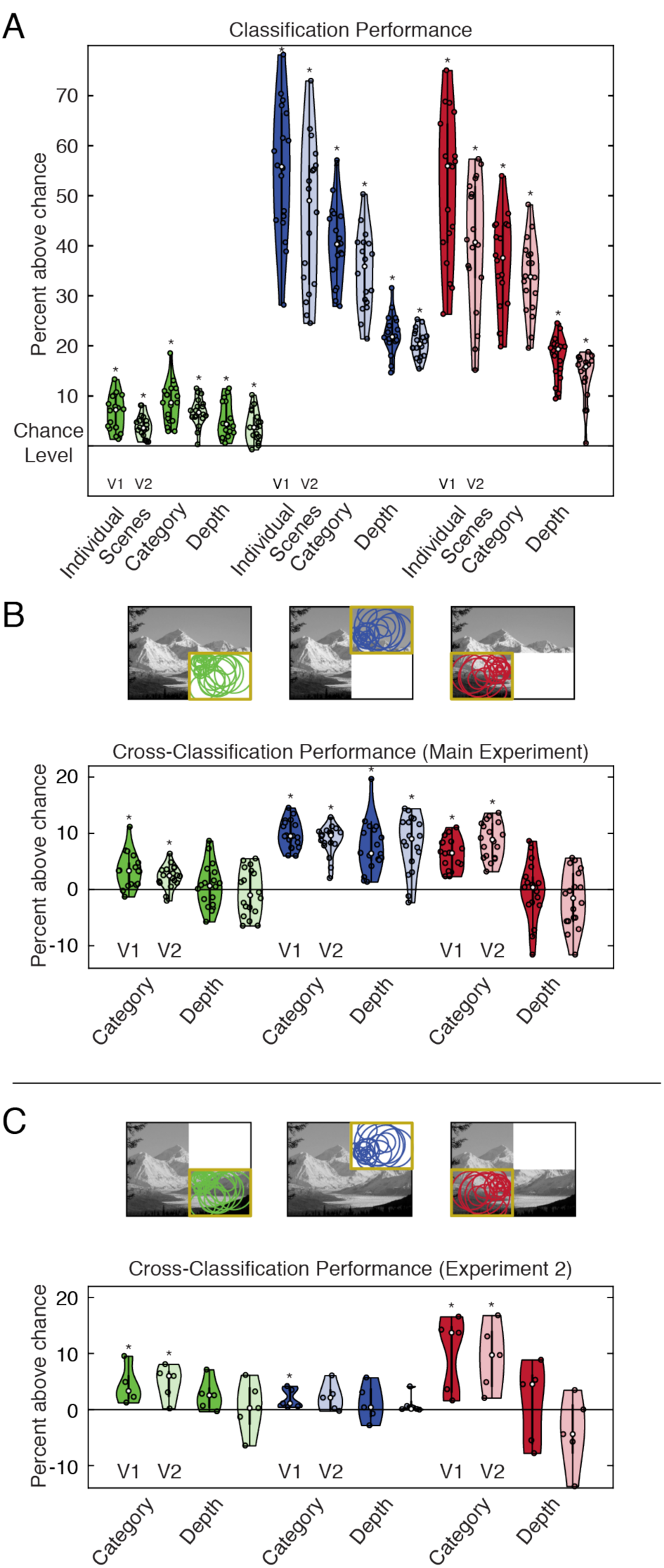
Classification and cross-classification performance. (A) Average classifier performances (N = 18) are shown for each visual area. Lower-Right, Upper-Right and Lower-Left quadrant analyses are shown in green, blue and red, respectively (the occluder covered the Lower-Right quadrant in the main experiment and the Upper-Right quadrant in Experiment 2). Individual subject data are shown as dots, and the distribution of these data were computed using kernel density estimation. Asterisks indicate greater than chance-level decoding accuracy (p < 0.05, one-sided Wilcoxon Sign-Rank test). Chance level is 4.17% for individual scenes, 16.67% for categories, and 50% for depth. (B-C) Cross-classification performance for Experiments 1 and 2 (N = 18 in Experiment 1; N = 5 in Experiment 2). Training occurred on 18 and 22 [of 24] randomly chosen scenes in category and depth analyses, respectively, and testing occurred on scenes not used for classifier training. Results were averaged over 100 iterations per subject.

To address our main question, we asked whether information was decodable from Occluded V1 and V2, which receive only cortical feedback and lateral interactions but no direct feedforward input. We were able to decode individual scene, category and depth information in V1 and V2 (Figure 2a), suggesting that internal models delivered to early visual cortex contain these types of information.

Our decoding results were also reliable at the level of individual subjects (Table S1; Figure S1); scene, category, and depth decoding were above chance-level in at least 14 of 18 subjects in nearly all regions tested. We found the weakest decoding for Depth in Occluded areas, which were only above chance-level in 10 (Occluded V1) and 7 (Occluded V2) of 18 subjects.

To further test whether Occluded V1 and V2 represent higher-level properties of scenes, we performed cross-classification analyses for scene category and depth information. We trained SVM models using responses to a subset of our scenes, leaving out a test-set for later cross-classification. For the category analysis, 18 (of 24) scenes were selected, leaving out one scene per category. For depth, we selected 22 (of 24) scenes, leaving out one scene per depth. We tested the classifier on the left-out scenes in a cross-classification approach. Due to the large number of possible image permutations in these analyses, we randomly assigned scenes to training and testing sets 100 times in each subject. Cross-classification of category was successful in Occluded and Non-Occluded areas. Cross-classification of depth was only successful in the Non-Occluded Upper-Right quadrant, suggesting that depth information is not available in lower visual field responses, regardless of whether feedforward information is available (Figure 2, Table S2).

Our cross-classification results show that responses in both Occluded and Non-Occluded areas of the lower visual field do not contain depth information. This visual field bias limits our ability to assess whether depth information is present in feedback to early visual cortex. We therefore conducted a second fMRI experiment in five subjects using the same scenes, but with the occluder moved to the upper-right quadrant of the visual field. In this experiment, we successfully cross-classified category in V1, with V2 not reaching significance (p = 0.091; one-sided Wilcoxon Sign-Rank). Once again, we were not able to cross-classify depth information in the Occluded quadrant (Figure 2). We therefore conclude that internal models of scenes contain category information but not depth information. This finding provides neuroscientific evidence that supports hierarchical views of visual processing. These views suggest that category information is useful for defining global context, which can then be fed back to the early visual cortex to aid with processing, for example in object recognition^13,14,19,20^.

### Decoding performance in V1 and V2

Differences in Occluded and Non-Occluded V1 and V2 decoding levels are informative for understanding how these areas might make distinct functional contributions to contextual scene features during feedforward and feedback processing. We found decoding to always be higher for feedforward (Non-Occluded) than for feedback (Occluded) conditions in both V1 and V2. Feedforward decoding (Non-Occluded) was higher in V1 than in V2 in both ROIs (upper and lower ROI) for individual scenes and category information. For depth decoding, feedforward decoding was higher in V1 than in V2 in the upper-right ROI but not in the lower-left ROI. For cortical feedback, there was no significant difference in the decoding of category or depth between V1 and V2, but there was a significant difference when decoding individual scenes. This difference between Occluded V1 and V2 decoding cannot be explained by ROI size, as voxel counts did not significantly differ (p = 0.116, one-sided paired t-test; see Figure S2). These findings support previous work suggesting that scene-specific features are more easily read out from V1 than from V2^4^. The current study’s larger scene set provides additional information, showing that more-general scene features relating to scene category and depth can be read out from Occluded V1 and V2 with approximately equal fidelity.

Visualising retinotopic information patterns

Our Occluded V1 and V2 decoding results detect reliable differences in response patterns between scene categories (but less so for depth), but the results do not indicate which scene features elicited these patterns. This consideration is particularly important in Occluded regions, where we would like to ensure that successful decoding is not simply due to fMRI signal spill-over or edge effects carried by lateral connections between Occluded and Non-Occluded areas at the occlusion boundary. To visualise classification information over corresponding scene features, we projected voxel classifier weights (the relative contribution of each voxel to a Linear Discriminant Analysis solution) into visual space (Figure 3). Positive weights indicate voxel responses associated with a classifier choosing the current scene and negative weights indicate responses associated with a classifier choosing a different scene. To assess whether weights were significantly different from zero, we performed two-sided t-tests across subjects at every pixel location in the visual field, and maps in Figure 3 show t-values surpassing a p < 0.05 threshold.

**Figure 3.**
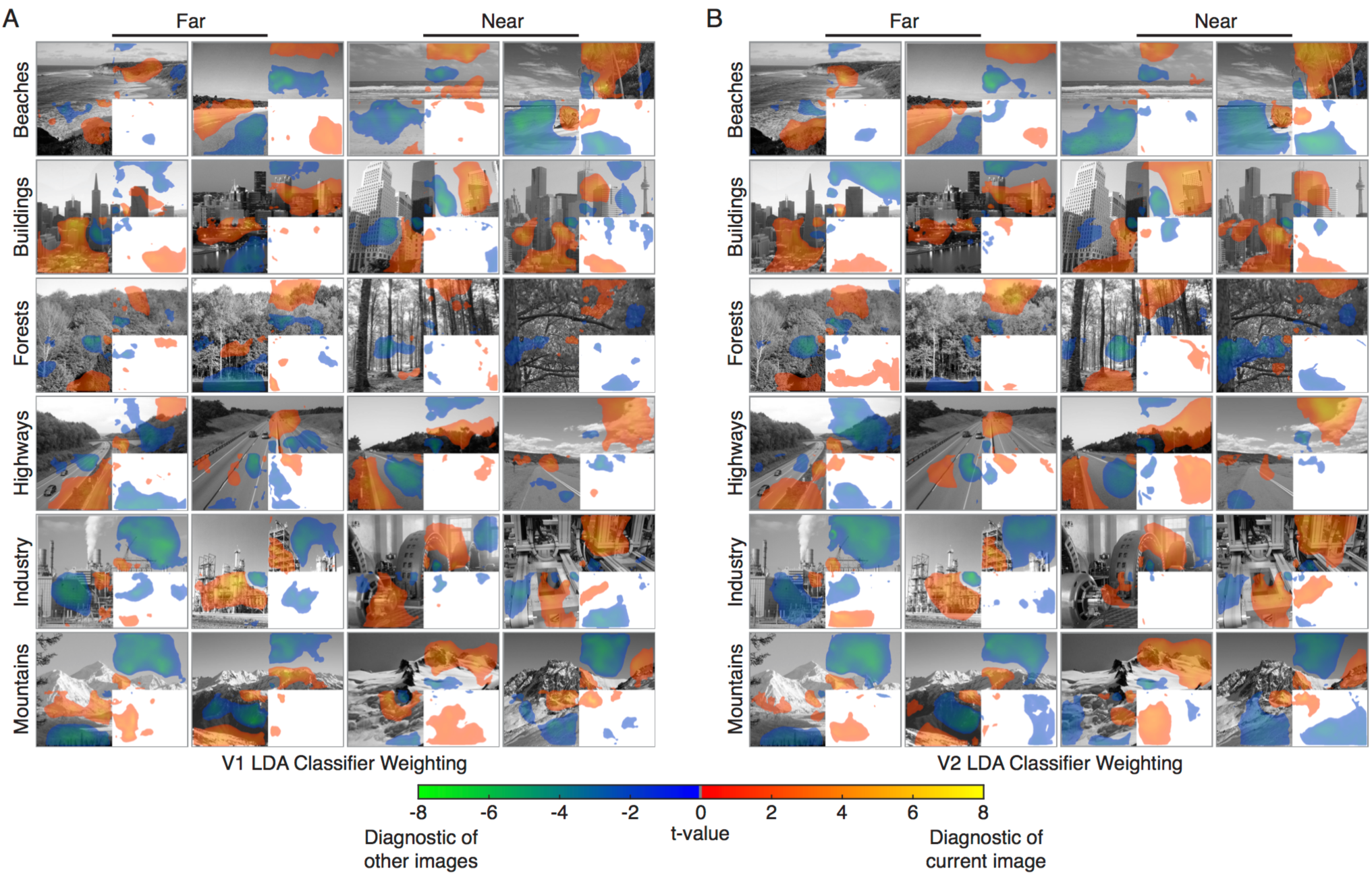
Projections of (A) V1 and (B) V2 Linear Discriminant classifier weights into visual space. Voxel classifier weights (the relative contribution of each voxel to an LDA solution) were averaged for significant classifications involving each scene, resulting in one map of discriminatory visual information for each scene in each subject. A two-tailed t-test was conducted across subjects at each pixel location in visual space to obtain t-value maps (p<0.05 threshold). Warm colors (red, orange and yellow) indicate visual areas of the scene where voxel activation is indicative of the respective scene. Cool colors (green and blue) indicate areas where voxel activation is indicative of a different scene.

Classifier information closely corresponds to visible scene features in Non-Occluded areas (Figure 3). Visual inspection shows that positive weights often match the edges of scenes or high-contrast scene areas and that negative weights match low-contrast areas, such as sky or water. Additionally, there is significant classifier information in Occluded areas despite relatively small response amplitudes in comparison to Non-Occluded areas (see Figure S3 for response amplitude maps). Crucially, Occluded classifier information does not reside along the Occlusion boundary. This result indicates that the signals we are examining are indeed from cortical feedback and are not due to signal over-spill or to edge effects produced by lateral interactions. In addition, responses do not continuously decay from the occlusion boundary, indicating that the recorded V1 and V2 responses are not due to lateral interactions spreading out from neighbouring compartments in the same cortical area. Instead, these responses could be explained by feedback signals alone, or by interactions between cortical feedback and lateral processing^21^.

### Line drawings as internal model read outs

We have shown that internal models transmitted to early visual cortex by cortical feedback convey scene category information. We also found that responses related to internal models could be used to decode individual scenes and that visual projections of classification information allowed us to see their retinotopic locations. Next, we wanted to understand whether internal models carry information about predictable occluded features. To derive these features, we conducted a behavioural experiment in which 47 participants drew in the missing scene information that they expected to be behind the white occluder. Figure 4 shows drawings generated by averaging across individual subject’s drawings. The coherency of individual drawings is apparent from visual inspection, emphasising the consistency of internal models across individuals. For some visual scenes, subjects’ drawings included most of the hidden scene features (well-predicted scenes), while other scenes had some of the features missing (poorly predicted scenes). Interestingly, the missing features were also consistent across subjects’ drawings. Representative examples of well-predicted and poorly predicted scenes are shown in Figure 4B, which compares drawings to hidden subsections of scenes. Drawings are highly consistent between individuals, leading us to conclude that their internal models of missing scene information are also highly consistent.

**Figure 4.**
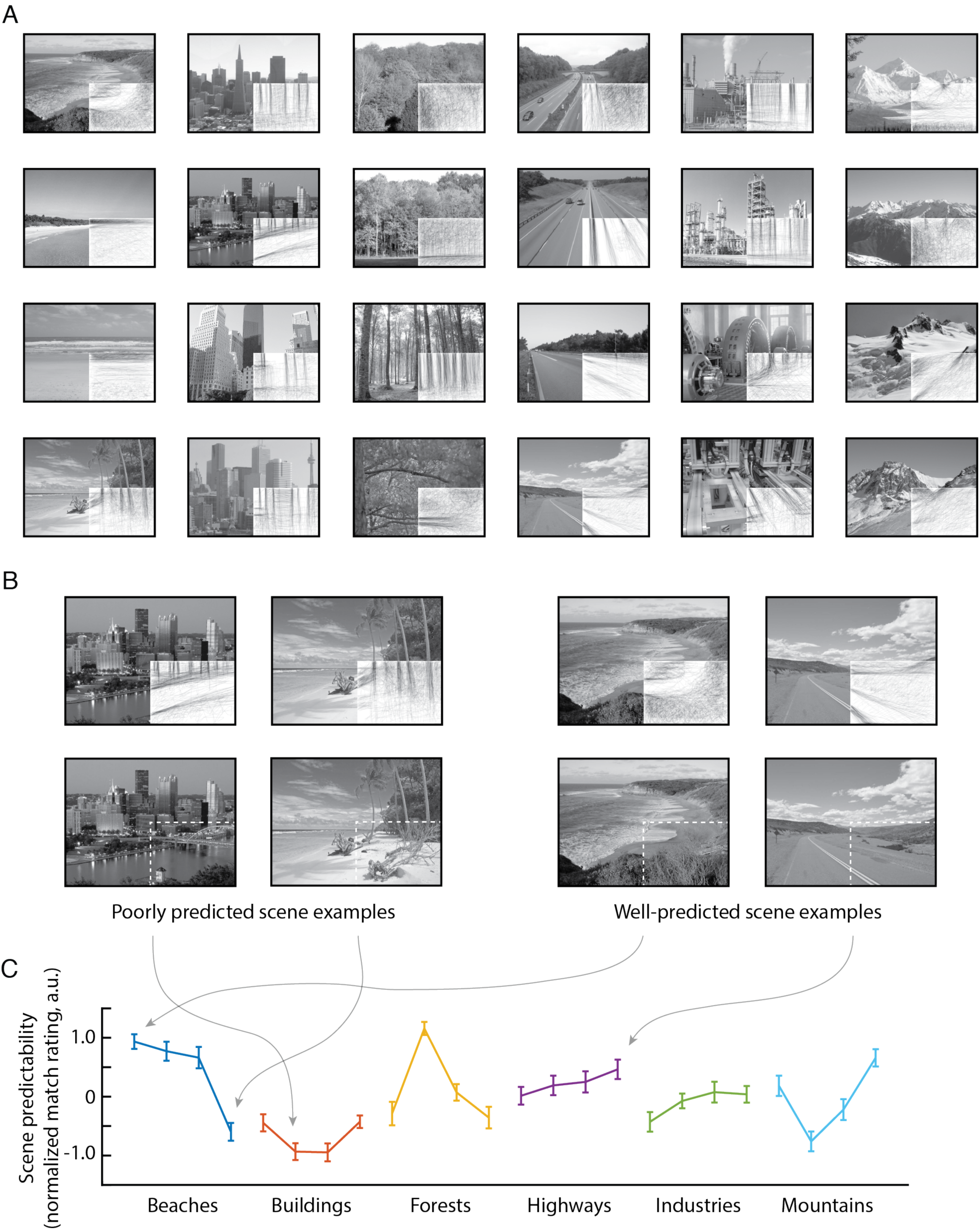
Line drawings are a behavioural measure of internal scene models. Forty-seven individuals filled in the occluded subsections of each scene as a line drawing. (A) Individual subjects’ drawings were averaged to create a single group-level drawing for each scene. (B) Hidden subsections of example scenes are compared with their respective line drawings. In both well-predicted and poorly predicted scenes, drawings are highly consistent across individuals. (C) A separate behavioural survey provided ratings of the predictability of each scene’s occluded section (organised by scene category). Ratings depict how well each drawing matched the actual missing portion of the scene. Mean and standard error of normalised values are displayed, and arrows highlight the example scenes from (B).

Using line drawings as behaviourally defined predictions, we used three visual feature models to predict scene representations in V1 and V2: the Weibull model, which corresponds to lateral geniculate contrast processing^22,23^; the Gist algorithm, which is similar to the orientation and spatial-frequency filters in V1^11^; and the H-Max model (Layer C2), which is matched to the tuning properties of intermediate ventral stream areas, such as V4 or posterior IT^20^. We computed these models on the line drawings and related them to brain activity in Occluded areas. To describe brain activity in Non-Occluded areas, we computed these three models (Weibull, Gist and H-Max) on the full greyscale scene data (from Upper-Right and Lower-Left quadrants). In addition to these scene-specific models, we also included a Category model and a Depth model. The Category model consisted of the 6 scene categories (Beaches, Buildings, Forests, Highways, Industries and Mountains). The Depth model was determined by a behavioural experiment in which 10 participants estimated the depth of each scene in meters (Figure S4).

Using Representational Similarity Analysis (RSA)^24^, we characterised the multivariate information similarity between scenes in our models and brain data. This allowed us to infer the information content of cortical feedback sent to Occluded brain regions. Figure 5 displays the similarity between models and brain data. In Occluded V1 (Figure 5A, upper panel), the Weibull, Gist and Category models are all significantly correlated with internal models from cortical feedback (p < 0.001, one-sided Wilcoxon signed-rank). Interestingly, the orientation information in line drawings (from the Gist model) is significantly more similar to Occluded V1 than Category is (p = 0.048, two-sided Wilcoxon signed-rank). This finding indicates that internal models delivered to Occluded V1 by cortical feedback are better described by orientation information from line drawings than they are by Category.

**Figure 5.**
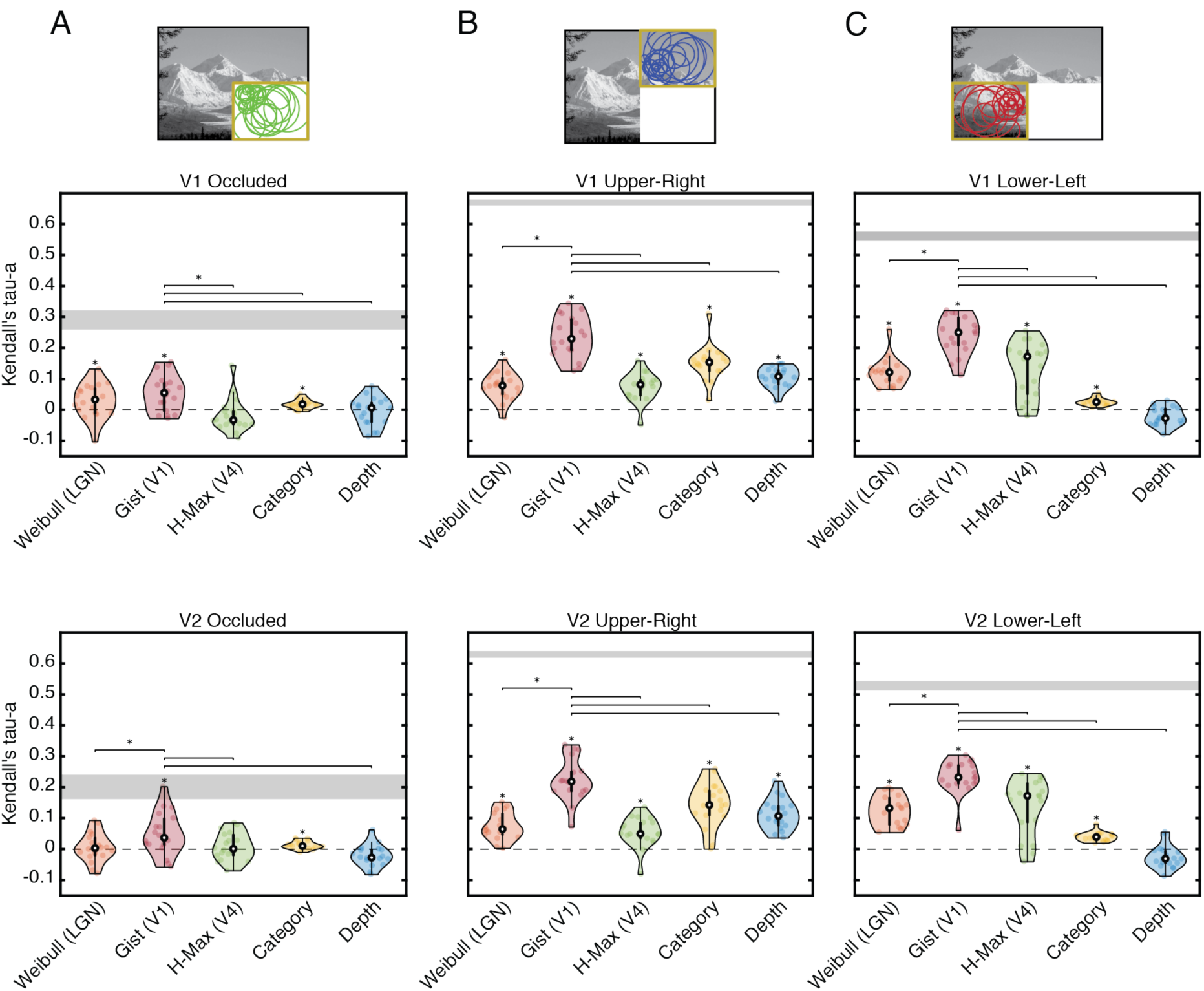
Comparison of scene-specific and global models with cortical representations. The similarity of each model with cortical representations in (A) Occluded and (B-C) Non-Occluded quadrants of V1 and V2 is shown as a rank correlation (Kendall’s Tau-a). The Weibull (LGN contrast processing), Gist (orientation and spatial-frequency processing) and H-Max (mid-level visual feature processing) model features were computed from visible scenes for Non-Occluded areas, and from line drawings for Occluded areas. Global feature models included Category and Depth (depth measurements were determined in a separate behavioural experiment). Individual subject data are shown as dots, and data distributions were computed using kernel density estimation. Asterisks directly above data indicate significantly greater than zero correlations (p < 0.05, one-sided Wilcoxon Sign-Rank test). Lines between models indicate significant differences in performance between the Gist model (the best performing model in all areas) and other models (see Figure S5 for all comparisons). Noise ceilings were calculated as the upper and lower bounds on individual subject correlations with an average representational structure.

In Occluded V2 (Figure 5A, lower panel), the Gist and Category models are significantly correlated with internal models from feedback. Here again, the Gist model is the highest-correlated model (higher than both Weibull and H-Max models [p = 0.039 and 0.025, respectively], but not higher than Category [p = 0.071]). In Occluded V2, individual subject correlations of the Gist model with feedback reach into the noise ceiling, indicating that orientation information in line drawings is a good model description for this dataset.

In Non-Occluded V1 and V2, all models are significantly correlated with brain data other than the Depth model in the Lower-Left quadrant. The Gist model performs significantly better than any other model (p < 0.01) in both Non-Occluded V1 and V2 (Figure 5B, C). This corresponds to the known language of early visual cortex, which is thought to respond to orientation information from feedforward visual input. In this context, our results reveal that cortical feedback to V1 and V2 is also translated into this “orientation language”.

### Internal models are generalised in predictable scenes

Our results show that early visual cortex responds to visual information hidden from view and that these responses are well-described by orientation information from line drawings. We have also shown that this behavioural readout of internal models is largely consistent across individuals, regardless of whether they accurately describe the hidden features of scenes (Figure 4B-C). To assess whether line drawings provide an improved model of feedback to V1 and V2, compared to the actual hidden scenes, we compared the performance of the Gist model computed from these two inputs. We found that the features derived from line drawings had statistically higher correlations with Occluded activity patterns than the features derived from actual scenes in V2 (p = 0.035, two-sided Wilcoxon Sign-Rank test across subjects), but not in V1 (p = 0.25). This finding implies that cortical feedback to V2 contains more generalised information than feedback to V1, since line drawings preserve the visual information required for scene interpretation, while disregarding other information^16,17^.

However, as previously noted, line drawings matched occluded scene features to varying degrees within the scene set. We measured this variability in a behavioural experiment in which 27 individuals rated how well line drawings and full scenes matched in side-by-side comparisons. Ratings can be interpreted as the predictability of the features hidden by the visual scene occluder (Figure 4C). To understand whether a relationship exists between scene predictability and model performance, we ordered scenes from most predictable to least predictable and repeated our previously described RSA modelling in bins of 8 scenes in a sliding-window fashion. Figure 6A shows how the Gist model performed when computed from line drawings and actual occluded scenes. Interestingly, when scenes are more predictable, models computed from line drawings outperform models computed from actual hidden scenes in both V1 and V2 (p < 0.05, two-sided Wilcoxon Sign-Rank test within each bin). Model performances converge as scenes become less predictable. This finding suggests that the internal models in cortical feedback to both V1 and V2 do indeed contain generalised scene information when scenes are easily predicted. However, line-drawing models are less advantageous compared to actual hidden scene information when the occluded portion is less predictable. In this case, subjects’ internal models might be more variable and thus not as generalised as they are when the occluded area is more predictable.

**Figure 6.**
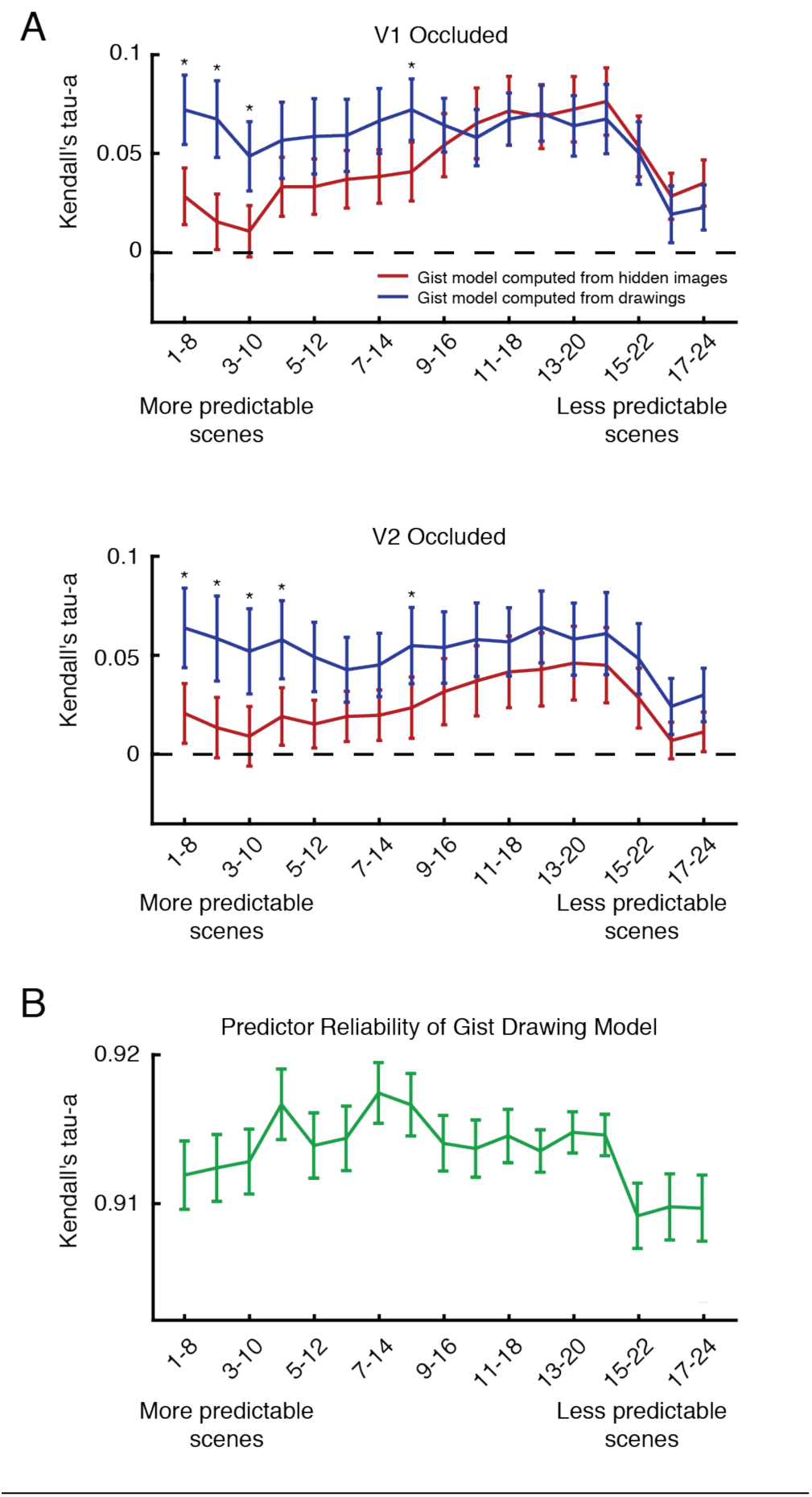
Relating scene predictability to models computed from line drawings and occluded scenes. (A) Occluded V1 and V2 correlations with Gist features computed from line drawings or actual hidden scenes are shown. RSA model comparisons were conducted in bins of 8 scenes, which were organised by scene predictability (see Figure 4C). Mean and standard error or correlation values are shown for each model and asterisks indicate significant differences between performances of the two models (p < 0.05, Wilcoxon Sign-Rank test). (B) The reliability of Gist features computed from line drawings are shown within each scene bin (mean and standard error).

In both V1 and V2, line drawing and actual hidden scene model performances both decrease in the least predictable scenes of our scene set. To see if this feature of our model predictions was related to behaviour, we performed a split-half reliability analysis on our Gist model features computed from line drawings (Figure 6B). We randomly split our subject group in half 50 times, averaged over individuals’ drawings in each iteration (resulting in two average drawings of each scene), computed Gist features and calculated the correlation between split-halves. While orientation information from the Gist model was highly reliable in all scenes (correlation values greater than 0.9), there was a distinct decrease in this reliability measure in the least predictable scenes, similar to model performances. Line drawings thus not only serve as a behavioural readout of internal models sent to V1 and V2, but their reliability also corresponds to our ability to predict cortical feedback patterns in these areas.

Overall, our results corroborate previous work showing that internal models conveyed by cortical feedback contribute to responses of visually occluded brain areas^3,4,6^. Here, we have extended our understanding of cortical feedback by showing that internal models sent to V1 and V2 contain category and scene-specific orientation information. Further, we discovered that responses associated with cortical feedback to V1 and V2 correspond with behavioural predictions of missing visual scene information in the form of line drawings. These results not only show that the earliest stages of cortical sensory processing are informed by behaviourally relevant predictions about the structure of the world, but also demonstrate that line drawings, an ancient method of conveying visual information, provide a means of accessing the brain’s internal models.

## Discussion

Our findings uncover activation patterns in occluded sub-regions of early visual cortex that were informative for determining category and individual scene information about the surrounding images. These results indicate that contextual feedback to early visual cortex exhibits high-level structure yet is scene-specific. We also found that this scene-specific information in feedback correlates with orientation information found in internal models of scenes, which we sampled by asking subjects to complete line drawings of the occluded subsections of scenes. Line drawings have remained largely unchanged during human history and therefore might embody fundamental components of how our visual systems represent the world^16,17^. Our findings support this hypothesis, particularly when scenes are highly predictable, where we have shown that line drawings outperform actual hidden scene features at predicting brain activity in occluded regions. This also supports a view of the visual system as a hierarchical inference network, with V1 acting as a geometric buffer or blackboard^25,26^. In other words, V1 preserves scene information for reference in calculations where image details or spatial precision are required.

Our results, and those of several other studies, support the idea that expected information is present in the early visual cortex without direct visual stimulation^3,6,25,27,28^. The expectation of visual features that are absent can lead to an illusory percept termed *modal completion*^7,29,30^. However, modal completion differs from the phenomenon we have investigated here. In this experiment, no conscious percept was triggered by the occlusion of visual features. Instead, we anticipate that knowledge captured in internal models supports *amodal completion* only, whereby a rudimentary expectation exists about the physical continuation of scene elements into the occluded portion of each scene, without a perceptual filling-in of those scene features.

Nevertheless, we are left with the question of how the relatively small effects that we observe in Occluded areas impact cortical processing. We hypothesise that cortical feedback signals play an important role in determining the output of neurons, and that this mechanism is linked to perception^31^. Evidence from animal models suggests that cortical layer 5 pyramidal cell spiking is virtually unaffected by stimulation of the apical tuft dendrites alone^32,33^, where cortical feedback is largely received. However, cells are tuned to be highly sensitive to associative feedback upon receiving feedforward input to their somatic dendrites in a process termed backpropagation-activated Ca2+ spike firing (BAC firing)^32^, where the coincident arrival of feedforward and feedback tuft inputs leads to bursts of action potentials. Since the BOLD signal in fMRI is sensitive to energy consumption in both dendritic synaptic processes and spiking activity, as has been shown by primate data^34,35^, we might be detecting in this study dendritic stimulation without feedforward input in Occluded regions of cortex. These synaptically driven BOLD responses should appear to be comparatively weak to those caused by the rigorous BAC firing that occurs when feedforward and feedback inputs are integrated. When combined, these points provide one explanation for how relatively small BOLD changes in Occluded regions can be associated with neuronal processes that significantly affect cortical processing prior to perception.

We computed visual feature models on our behavioural line drawings and related them to brain activity in regions receiving cortical feedback signals. By describing which model fits best to brain activity, we have enhanced our understanding of how internal models are communicated in a hierarchical visual system. One assumption is that the top-down stream takes information from a feature space in a hierarchically higher-level visual area and translates it to a feature space used in a lower-level visual area. This idea of inheriting the features of a higher processing stage has been shown in the hierarchically organised macaque face processing network^36,37^. For example, the tested H-Max model (C2 level) explains a V4-or Posterior IT-like processing stage in a feedforward network^20,24^. Consequently, using RSA we found significant correlations with visual responses in Non-Occluded areas that process feedforward information. However, the H-Max features computed from behavioural line drawings did not correlate with brain activity carried by cortical feedback (i.e. in Occluded area responses). This finding indicates that the mid-level features computed by an H-Max-C2 model might describe the feedforward feature space, but that cortical feedback represents different features. Hence, the mid-level features summarised by the H-Max model might not contribute essential aspects of the internal models that are transmitted to early visual cortex. In contrast, the lower-level visual features of the Gist model correlated with both Non-Occluded and Occluded responses, indicating that this low-level feature space is a common language used by both feedforward and feedback processing. Orientation and spatial frequency, which are calculated by the Gist model, might be important for predictions in V1 and V2. However, after modelling the behaviour tested by line-drawings, we conclude that the optimal models for describing cortical feedback responses are those that use less features, namely those expressed in line-drawings and in the Gist model.

We found that depth information was not present in feedback to V1 and V2. It is important to note that Non-Occluded V1 and V2 responses in the lower visual field also did not contain depth information, at least for the current stimuli set. This finding highlighted the possibility that our task missed depth information in feedback that was arriving at the upper visual field. We therefore conducted a second fMRI experiment with an upper visual field occluder. This second study replicated the initial finding – depth information was again not present in occluded activity (now in the upper visual field), and the non-occluded lower visual field also did not contain depth information. Future studies would therefore need greater sensitivity to test whether scene depth forms part of cortical feedback information and its contribution to processing biases in V1 and V2 for upper versus lower visual field. A large-scale study has recently reported that several localised early visual cortical patches respond preferentially to objects nearer to the viewer^38^. The ability to systematically map depth properties onto the cortex while manipulating the availability of feedforward image information would enable researchers to resolve the contributions of feedforward and feedback signals to these cortical response properties.

We found that line drawings replicate the structure of internal models in early visual cortex. Importantly, drawings and brain activity recordings were from different groups of subjects. This provides evidence that line drawings contain generalisable features related to the structure of internal models across individuals but begs the question of whether individual differences exist in this relationship. In future, it will be important to carefully study line drawings from individuals that also have brain recordings like those from the current study to understand if differences in participants’ line drawings predict differences in the structure of their internal models.

Artificial neuronal networks have achieved high performance on a number of natural signal processing tasks including visual object and speech recognition^39^. However, such networks predominately use feedforward architectures, and therefore perform sub-optimally when presented with partially occluded visual scenes. Recent work has shown that by including feedback and lateral connections, convolutional networks outperform those with only feedforward connections on tasks involving occlusion^40^. In this study, we have shown that Occluded V1 and V2 responses contain predictive information about missing feedforward input and that this information can be modelled using behavioural drawings. Based on these results, we propose that feedback and lateral connections are important for models that aim to replicate human performance. Furthermore, line drawings can be incorporated into training sets as targets for non-feedforward connections within networks. Recent work was able to reproduce drawings of isolated common objects^41^, but has not used line drawings as a source of scene predictions. By incorporating behavioural sampling of human internal models using line drawings, artificial neuronal networks might be able to explain aspects of cortical responses not yet captured by current computational models and thus provide insight into the mechanisms of feedback processing in early visual cortex.

Our findings advance our understanding of the visual features conveyed by cortical feedback. The mental models expressed at the first processing stage in our visual cortex are closely related to mental models depicted in line drawings. When faced with a blank canvas, it is conceivable that an artist drawing a visual scene represents this scene using cortical feedback processing, until their line drawings converge with the internal model of the visual scene. In the present study, line drawings were consistent across individuals. This is presumably because human visual systems have a comparable mechanism for projecting internal models in a top-down manner to the cortical stage for visual information, with this stage acting as an active canvas or blackboard^25,26^.

Accessing the brain’s internal models is a key advancement that cognitive neuroscience will need to focus on in the next decade. Our current work concentrated on the visual system because it is well suited for reading out internal models – this is however only an example of how internal models contribute across the brain. Internal models play an active role in many human behaviours including visual attention^42^, higher cognitive functions such as memory and action planning^43^, decision making^44^ and mental time travel^45^. Moreover, internal models are affected in mental disorders including major depression^46^, schizophrenia^47^ and autism spectrum disorder^48,49^. Previous work has shown differences in the characteristics of line drawings made by schizophrenic patients^50^ and in the ability of autistic children to predict objects from fragmented line drawings^51^. In the context of our results, these studies might have been assessing differences in the structure of internal models in schizophrenia and autism.

By continuing to expand our brain reading of internal models, we can gain new insights into fundamental neuroscientific questions in health and disease.

## Methods

### Participants

Twenty-three healthy individuals (N = 18 in the main fMRI experiment: 12 female, age = 26.45 ± 5.70, mean ± SD; N = 5 in the second fMRI experiment: 2 female, age = 26.50 ± 5.58) with normal or corrected-to-normal vision gave written informed consent to participate in this study, in accordance with the institutional guidelines of the local ethics committee of the College of Science & Engineering at the University of Glasgow (#CSE01127).

### Stimuli

Twenty-four real-world scenes from six categories were chosen from a previously compiled dataset^9^. Images were displayed in grey-scale (matched for global luminance) on a rear-projection screen using a projector system (1024 × 768 resolution, 60 Hz refresh rate). Stimuli spanned 19.5 × 14.7° of visual angle and were presented with the lower-right quadrant occluded by a white box (occluded region spanned ≈ 9 × 7°). A centralised fixation checkerboard (9 × 9 pixels) marked the centre of the scene images. Stimuli in Experiment 2 were identical to this, with the exception that the occluder was moved to the upper-right quadrant.

### Experimental Design

Each of the eight runs consisted of six blocks of eight sequences of stimulation with intervening fixation periods, plus two mapping blocks (total scanning time per run was 804s). Each stimulation sequence lasted 120s, with 12s fixation at the beginning and the end of each series. Stimuli were flashed at a rate of 5Hz in order to maximise the signal-to-noise ratio of the BOLD response^52^. Each sequence was presented in a pseudo-randomised order where individual images were not shown repeatedly. Over the course of the experiment, each scene was presented 16 times (two times per run). To ensure fixation, we instructed participants to respond via a button press to a temporally random fixation colour change. So that participants would attend to the scenes, participants were asked to report the category of the scene being presented during the fixation colour change using 6 randomised response buttons.

We used mapping blocks to localise the cortical representation of the occluded region^7^. In a block design, subjects viewed contrast-reversing checkerboard stimuli (5Hz) at three visual locations: Target (lower-right quadrant in main experiment, upper-right quadrant in Experiment 2), Surround (of the target), and Control (remaining three quadrants). Each condition was displayed for 12s with a 12s fixation period following, and mapping blocks were randomly inserted between experimental blocks, once per run. We conducted retinotopic mapping (polar-angle and eccentricity) runs separately from the main experiment.

### fMRI Acquisition

fMRI data were collected at the Centre for Cognitive Neuroimaging, University of Glasgow. T1-weighted anatomical and echo-planar (EPI) images were acquired using a research-dedicated 3T Tim Trio MRI system (Siemens, Erlangen, Germany) with a 32-channel head coil and integrated parallel imaging techniques (IPAT factor: 2). Functional scanning used EPI sequences to acquire partial brain volumes aligned to maximise coverage of early visual areas (18 slices; voxel size: 3mm, isotropic; 0.3mm interslice gap; TR = 1000ms; TE = 30ms; matrix size = 70×64; FOV = 210×192mm). Four runs of the experimental task (804 vol.), one run of retinotopic mapping [session 1: polar angle (808 vol.); session 2: eccentricity (648 vol.)], and a high-resolution anatomical scan (3D MPRAGE, voxel size: 1mm, isotropic) were performed during each of two scanning sessions.

### fMRI Data Preprocessing

Functional data for each run were corrected for slice time and 3D motion, temporally filtered (high-pass filter with Fourier basis set [6 cycles], linearly detrended), and spatially normalised to Talairach space using Brain Voyager QX 2.8 (Brain Innovation, Maastricht, Netherlands). No spatial smoothing was performed. These functional data were then overlaid onto their respective anatomical data in the form of an inflated surface. Retinotopic mapping runs were used to define early visual areas V1 and V2 using linear cross-correlation of eight polar angle conditions.

A general linear model (GLM) with one predictor for each condition (Target > Surround; mapping conditions from experimental runs) was used to define regions of interest (ROI) that responded to the visual target region (lower-right quadrant) and two control regions (upper-right and lower-left quadrants), within V1 and V2. We then performed pRF analyses^18^ on all ROI voxels and excluded those voxels whose response profiles were not fully contained (within 2*σ* of their pRF center) by the respective visual ROI. Lastly, a conjunction of two GLM contrasts (Target > Surround & Target > Control for Occluded ROIs, and Control > Surround & Control > Target for Non-Occluded ROIs) was used to exclude any voxels responding to stimuli presentation outside their respective visual ROI (see Figure 1, Figure S2). Time courses from each selected vertex were then extracted independently per run and a GLM was applied to estimate response amplitudes on a single-block basis. The resulting beta weights estimated peak activation for each single block, assuming a standard 2*γ* hemodynamic response function.

### Classification Analyses

For SVM classification analyses, a separate regressor modelled each experimental trial. This procedure yielded a pattern of voxel activations for each single trial, and parameter estimates (*β* values) were obtained for each voxel and then z-scored. A Linear SVM classifier was trained to learn the mapping between a set of all available multivariate observations of brain activity and the particular scene presented, and the classifier was tested on an independent set of test data. Classification analyses were performed using a pairwise multiclass method. Classifier performance was assessed using an n-fold leave-one-run-out cross-validation procedure where models were built on [n – 1] runs and were tested on the independent nth run (repeated for the eight runs of the experiment). In analyses of category and depth-based classification, individual scene presentation labels were combined based on these distinctions before training and testing of the SVM classifiers. The significance of individual subject testing was assessed using permutation testing of SVM classifiers. We shuffled data labels in training sets and left testing set labels intact, repeating this procedure 1000 times. This procedure resulted in a null classification model around chance-level, and our observed classification value was compared to this distribution to determine the classification significance compared to chance. To determine the group-level distribution of classification performances, we averaged cross-validation folds of individual subject results to arrive at one performance metric for each subject per analysis. We then conducted non-parametric one-sided Wilcoxon Sign-Rank tests to determine whether the distribution of subject performances was above chance level.

Cross-classification analyses were performed similarly to those of our cross-validated classification, but our scene set was split up prior to model training. Training set sizes consisted of 18 and 22 scenes for category and depth analyses, respectively, and testing sets consisted of the remaining scenes. Due to the large number of possible scene permutations, we conducted 100 iterations of our analyses in each subject. For these analyses, training and testing sets were defined in a pseudo-random manner, where each category or depth was evenly represented within both sets. As in our cross-validated classification, training and testing of models occurred on independent data sets using a leave-one-run-out procedure. Since we performed leave-one-run-out cross-validation on each of the 100 training/testing sets, permutation testing for individual-subject classification significance was not feasible, as it would have required 100,000 tests in each ROI, information type and subject. We employed Wilcoxon rank-signed testing to examine individual subject performance. In each of the 100 training/testing sets, we averaged the 8 cross-validated classification performances, resulting in 100 performance values and tested those values against chance-level. To report group-level performance in these analyses, we averaged over cross-validation folds and then averaged performances over the 100 training/testing sets to get an individual subject performance. These values were subjected to one-sided Wilcoxon Sign-Rank tests to assess significance above chance.

### Retinotopic projections

Using each voxel’s two-dimensional Gaussian response, as estimated in our pRF analysis, we projected voxel classifier weights and activity patterns into visual space. Voxel pRF functions were multiplied by their respective weights from a Fisher Linear Discriminant Analysis^53^ to produce a single visual field map for each scene comparison. All comparisons involving a scene that had significant classification were averaged to produce a map for each scene in each subject. We then performed a two-sided t-test across subjects at every pixel location in the visual field in each scene to assess whether weights were significantly different from zero. This procedure was repeated with voxel responses to each scene (mean removed) to produce Figure S3.

### Line Drawings

Forty-seven individuals consented to participant in a behavioural experiment in which they filled in the occluded subsections of our scene set as line drawings. The experiment consisted of completing scenes using an electronic drawing pen and an Apple iPad Mini tablet (1^st^ Generation; screen resolution: 1024×768). The pen stroke was 6 pixels wide and produced purely black lines (no graded pressure settings). The tablet was held at approximately 45cm from the participant to approximate the visual angle of scenes in our fMRI experiments. Participants were given 25 seconds to complete each drawing with a 5 second break between drawings. The total length of the experiment was 12 minutes.

Line drawings were averaged over participants to capture the consistency of internal models. Therefore, lines drawn by a large proportion of subjects appear darker in Figure 4 than those drawn by a smaller proportion of subjects. Drawings were then scaled between 0 and 1 across the entire scene set, preserving the precision of internal models across scenes. Drawings were then used as input to visual processing models for Occluded ROIs.

To measure how well subjects’ line drawings captured the occluded portions of the test scenes, we conducted a behavioural experiment in which participants rated how well the line drawings and full scenes matched. Twenty-seven individuals were shown side-by-side comparisons of each scene in its non-occluded form and with its average line drawing in the occluded area. Scenes were presented in random orders and subjects rated the match on a scale from 1 to 7. Subjects were not given any time constraints, and the task took approximately 2-3 minutes. Ratings for the 24 scenes were z-scored within each subject and averaged across subjects to obtain a single predictability rating for each occluded scene.

### Model Comparisons

We compared visual processing models (Weibull, Gist, H-Max, Category and Depth; see individual model sections for detail) using an Representational Similarity Analysis framework^24^. In each visual ROI (visible portions of scenes in Non-Occluded regions and behavioural drawings or actual hidden scenes in Occluded regions), representations were calculated for individual channels of each model using a squared Euclidian distance metric (matrix sizes were scene comparisons [276] × model channels).

Model representational dissimilarity matrices (RDMs) were fit to data RDMs using non-negative least-squares^54^ in a cross-validated manner. First, two independent RDMs were calculated using the Linear Discriminant Contrast^53^ method (e.g. runs 1 & 2 vs. runs 3 & 4 and runs 5 & 6 vs. runs 7 & 8). For each scene comparison, models were fit to the RDM of the 22 other scenes in the first set (e.g. runs 1 & 2 vs. 3 & 4). This was repeated for all scene comparisons, thus producing a predicted RDM based on model parameters, which was then compared to the second half of the data (e.g. runs 5 & 6 vs. 7 & 8) using Kendall’s Tau-a rank correlation^55^. This procedure was repeated for all 70 possible split-quarter combinations, and values were averaged over splits to produce one correlation value per subject per ROI/model combination.

Noise ceilings for each cortical area were calculated as the upper and lower bounds on individual subject correlations with the average cortical RDM^55^. We measured the correlation (Kendall’s’ Tau-a) of each subject’s cortical RDM with the average subject RDM and defined the upper bound of the noise ceiling as the average of these correlation values. We repeated this procedure in a Leave-One-Subject-Out fashion to define the lower bound of the noise ceiling.

These analyses were repeated on subsets of scenes to understand whether there was a relationship between scene predictability and model performance of Gist features computed from line drawings and actual hidden scenes. We ordered the scene set from the most predictable to the least predictable (from our behavioural analysis of scene predictability, using ratings of how well line drawings matched hidden portions of scenes) and performed RSA modelling in 17 bins of 8 scenes in a sliding-window fashion.

### Weibull Model

The Weibull image contrast model measures the distribution of contrast values for an image and seeks to emulate the X and Y cells in the Lateral Geniculate^23^. It therefore had two outputs (Beta and Gamma statistics, corresponding to X and Y cells), which we calculated within each quadrant in areas extending from fixation to 1.5 and 5 degrees of visual angle, respectively^22,56^.

### Gist Model

The Gist algorithm measures the distribution of oriented bandpass Gabor filter responses in localised portions of images. Our model used default settings of 16 receptive fields (4 × 4 grid), 8 orientations, and 4 spatial frequencies^11^. This model had a 512-dimensional output.

We performed a split-half reliability analysis on Gist model features computed from line drawings to compare to model performances. We randomly split our subject group in half 50 times (23 subjects in one half and 24 subjects in the other half), averaged their drawings in each iteration (resulting in two drawings of each scene), computed Gist features and calculated the correlation between split-halves. We calculated the mean and standard error of scenes within each bin of 8 (organised by scene predictability).

### H-Max Model

The H-Max model is a hierarchical model that gradually combines visual features of higher complexity. Here, we used the output of its fourth sequential stage, C2. The first two stages (S1 and C1) correspond to the simple and complex cells or early visual cortex. Stages S2 and C2 use the same pooling mechanisms as S1 and C1, but pool from the C1 stage and respond most strongly to a particular prototype input pattern. Prototypes were learned from a database of natural images outside of this study^20^. The output of this model had 2000 dimensions.

### Category Model

Our Category model had six channels (one for each scene category). Comparisons between scenes within the same category took distance values of 0 and comparisons between scenes from different categories had distance values of 1.

### Continuous Depth Model

To quantify the depths of our scenes, 10 subjects were asked to assess the depth in meters of each scene in our scene set. To ensure we were probing the perceptual depth of each scene, participants were given minimal guidance on the definition of the term “scene depth.” Depth ratings were converted to a log_10_ scale, bootstrapped via 1000 samples of the mean, and a normal distribution was fit to the resampling histogram for each scene. This produced probability distributions for each scene’s depth^12^ (Figure S4). Squared differences between distribution means were used in our model comparisons analysis. This model had a single dimension as output.

## Acknowledgments

We would like to thank Luca Vizioli and Nikolaus Kriegeskorte for useful discussions regarding Representational Similarity Analysis, Fraser Smith for discussion of SVM classification analyses, and Steven Scholte for discussion and code for the Weibull model. We would also like to thank Matthew Bennett and Fiona McGruer for useful comments on the manuscript. This project has received funding from the European Union’s Horizon 2020 Programme for Research and Innovation under the Specific Grant Agreement No. 720270 and 785907 (Human Brain Project SGA1 and SGA2), and the European Research Council (ERC) under the European Union’s Horizon 2020 research and innovation programme under grant agreement No ERC StG 2012_311751-‘Brain reading of contextual feedback and predictions’ awarded to LM.

## Author Contributions

L.S.P. and L.M. conceptualised the study; A.T.M. and L.S.P. collected data; A.T.M. analysed data; A.T.M., L.S.P. and L.M. wrote the manuscript; L.M. acquired funding.

## Supplementary Information

**Figure S1.**
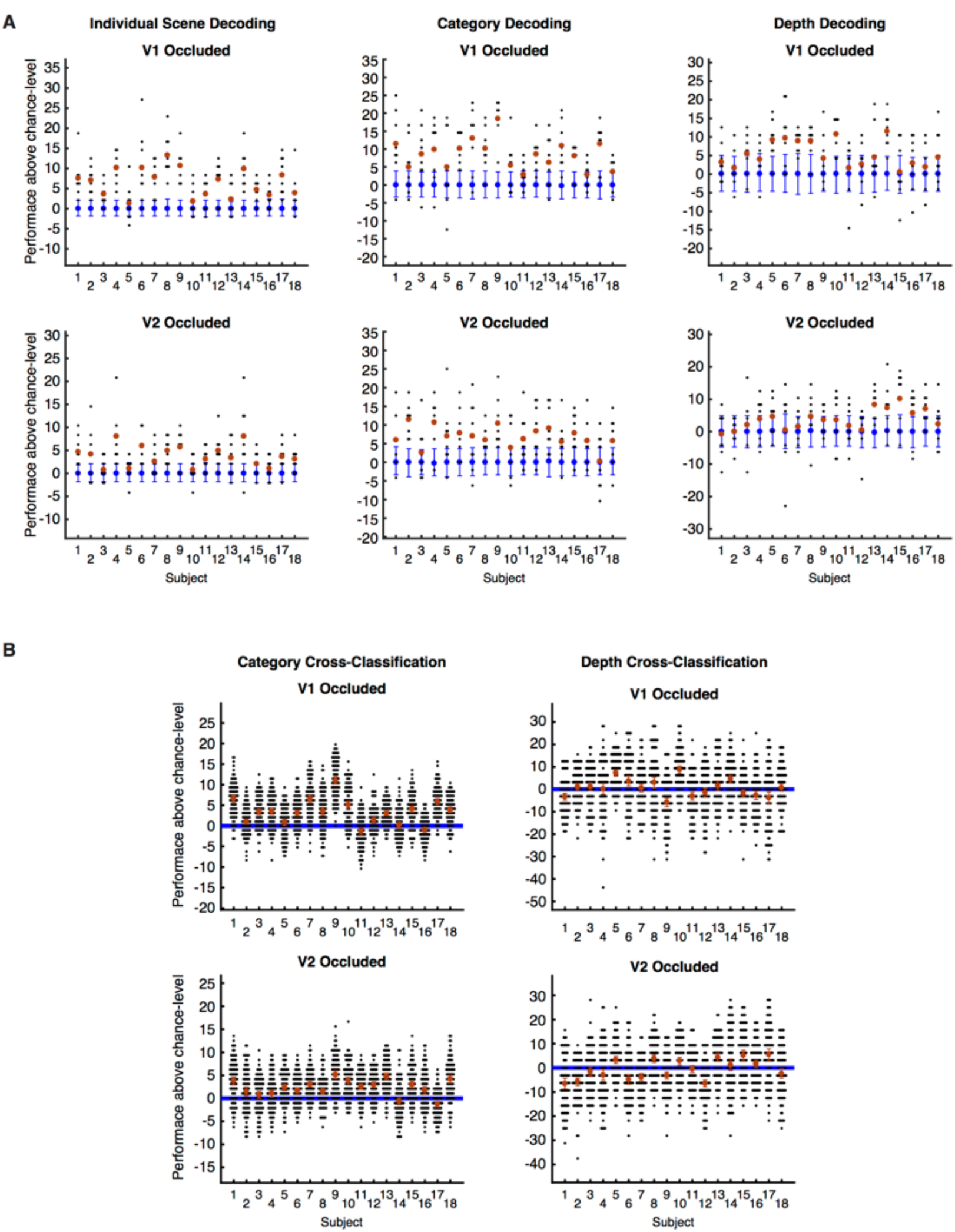
Individual results from SVM decoding analyses. (A) Classification performances compared to chance-level for each analysis type. Performances from each leave-one-run-out fold are shown as black dots, with the mean of all folds shown in red. 95% confidence intervals on a permutation-based null distribution are shown in blue. (B) Individual results from SVM cross-classification analyses. Individual performances on random splits of the scene set are shown as black dots, and 95% confidence intervals on the mean performance are shown in red. Chance-level performance is shown as a blue line.

**Figure S2.**
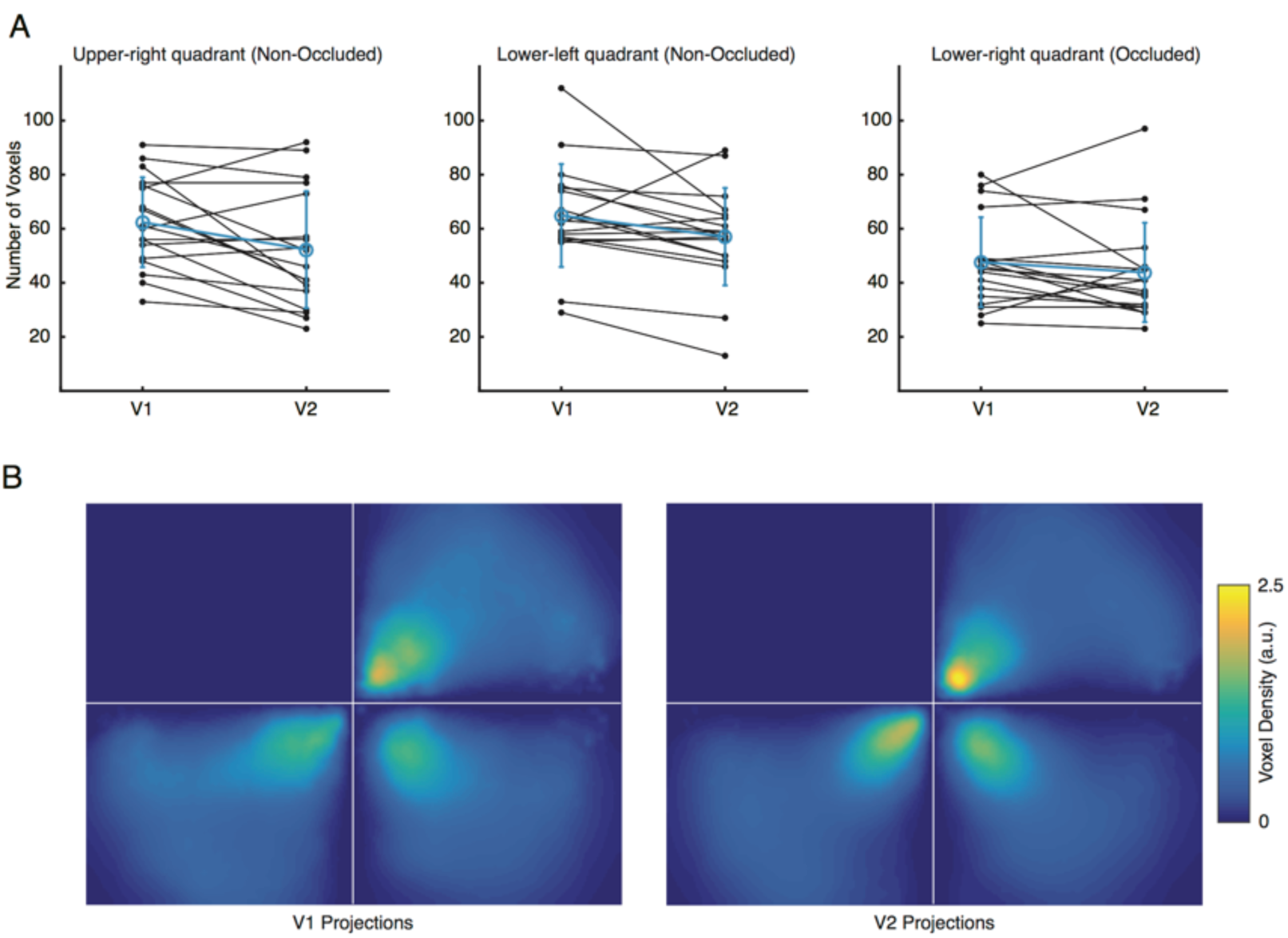
(A) V1 and V2 voxel counts in each region of interest. Black points and lines indicate individual subject voxel counts in V1 and V2 in each region of interest, and group statistics are shown in blue (with standard error). (B) Voxel density in regions of interest in visual space. Voxel density maps were calculated using the inverse of Scott’s Rule-of-Thumb (kernel width of 1° of visual angle) in individual subjects, masked by voxel coverage maps, and averaged across subjects. We tested whether the number of voxels in V1 was greater than the number in V2. There was a significant difference in voxel counts for Upper-Right and Lower-Left quadrants, but not in the Lower-Right Occluded quadrant (p = 0.007, 0.016, and 0.116, respectively, one-sided paired t-tests). Occluded areas were the only quadrant to not have significant differences in voxel counts between V1 and V2. We conclude that differences in classification analyses between Occluded V1 and V2 are not due to differences in voxel counts. We mapped the voxel density from our visual space analyses using the inverse of Scott’s Rule-of-Thumb for the bandwidth of a 2-dimensional kernel density estimator:

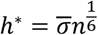

Where *h*∗ is the weighted average of pRF sizes based on a Gaussian kernel 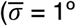 of visual angle), and *n* is the weighted number of voxels based on the same kernel. The kernel was convolved with the image and multiplied by a map of voxel coverage. Figure S1B shows the group average (N = 18) of individual subject maps.

**Figure S3.**
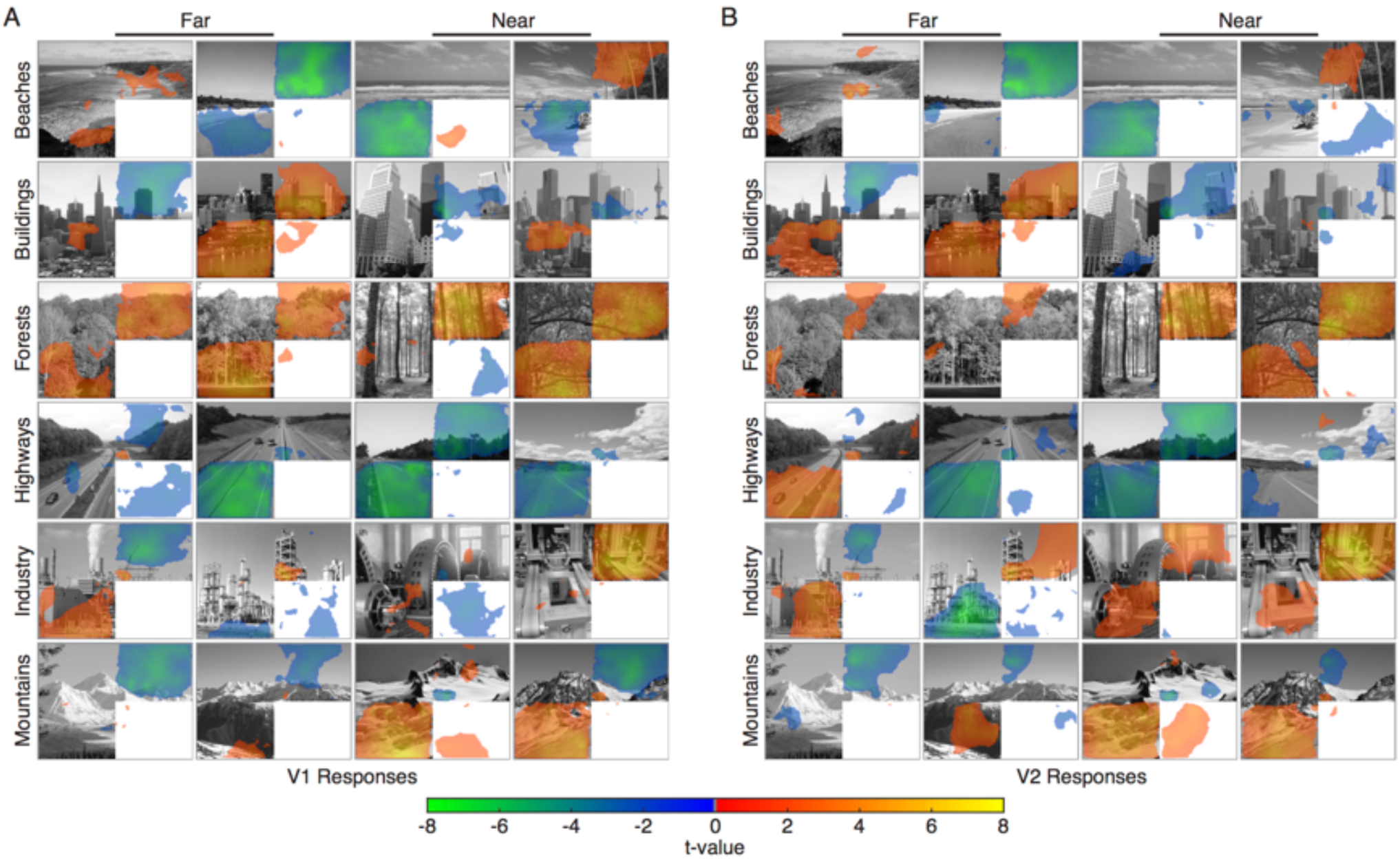
Projections of (A) V1 and (B) V2 response patterns into visual space. Voxel responses versus baseline were projected to visual space by calculating a weighted average of all voxels’ pRFs, where weights were each voxel’s response amplitude with the mean response to all scenes removed. Projections were mapped in individual subjects, and a two-tailed t-test was conducted across subjects at each pixel location in visual space to obtain t-value maps (p < 0.05 threshold). Warm colors (red to yellow) are above mean responses, and cool colors (blue to green) are below.

**Figure S4.**
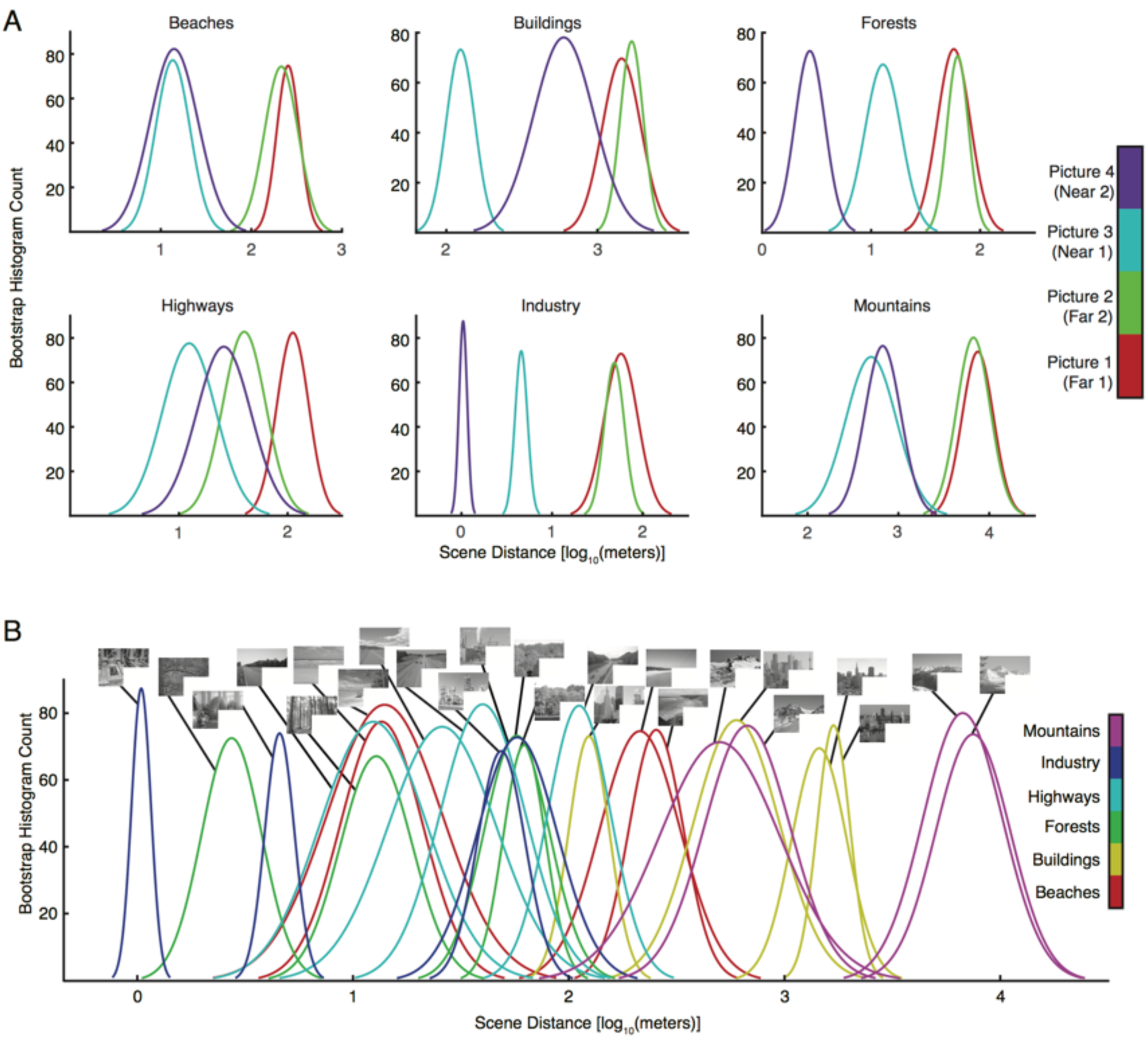
Behavioural depth ratings of scenes. (A) Probabilities of depth ratings (log_10_[meters]) for our 24 scenes are shown, grouped by category. Ratings were obtained via a behavioural experiment where subjects were asked to rate the depth of the overall scene in meters (N=10). Individual ratings were converted to log_10_(meters) and the normal distribution of mean ratings for each scene was calculated from a bootstrap histogram (1000 samples). (B) Depth ratings for our 24 scenes (identical to (A) but plotted together).

**Figure S5.**
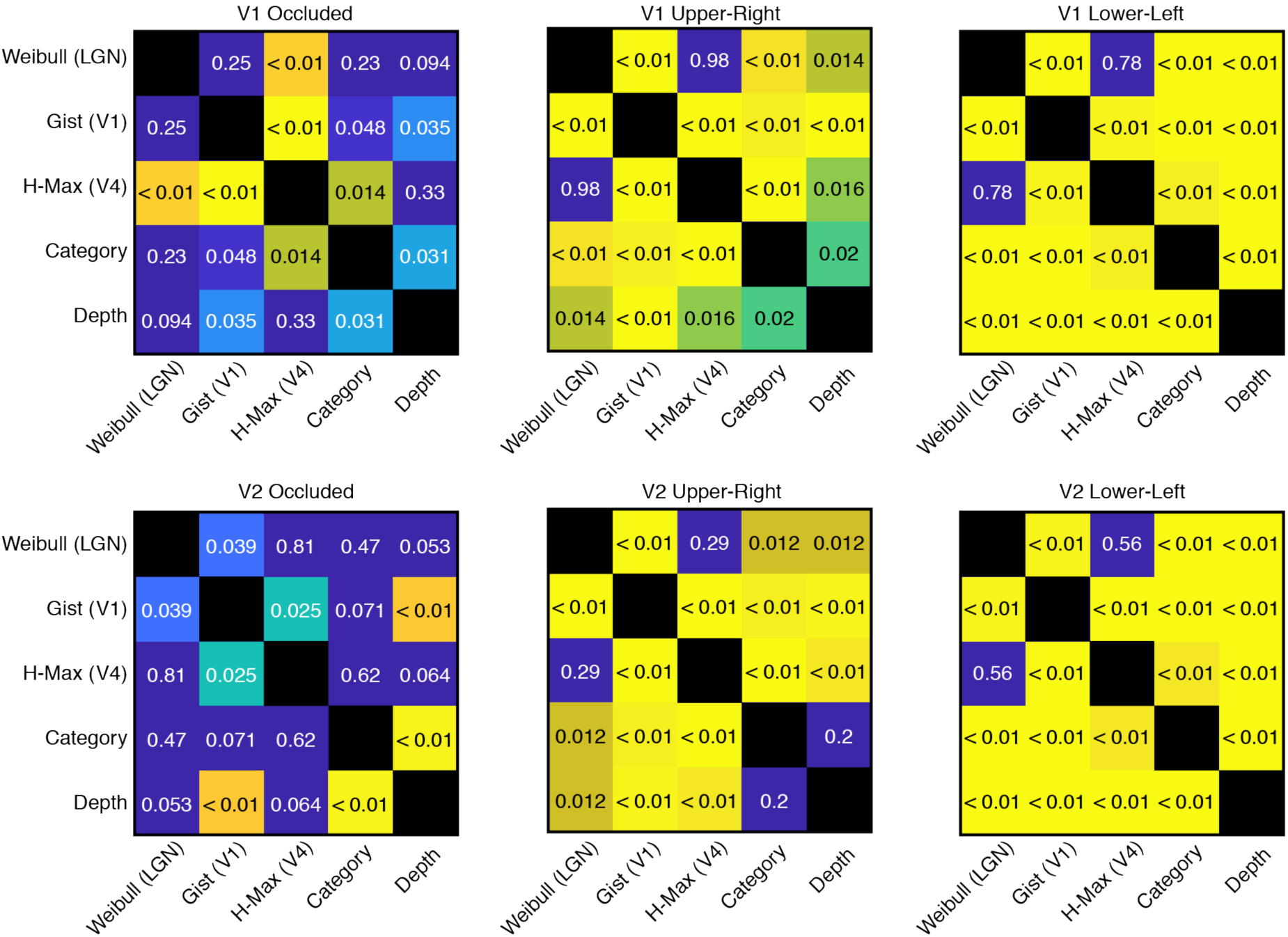
RSA model performance comparisons corresponding to Figure 5. Significance values (p-values from Wilcoxon Sign-Rank tests) for comparisons between model performances in Figure 5 are shown. Dark blue colors indicate comparisons not reaching significance. Significant comparisons appear as light blue, green and yellow, with yellow being most significant.

**Table S1.** Subjects with significant individual subject classifications (one-sided Wilcoxon Sign-Rank test).

|  | V1 |  |  | V2 |  |  |
| --- | --- | --- | --- | --- | --- | --- |
|  | Ind. Scenes | Category | Depth | Ind. Scenes | Category | Depth |
| Upper-right quadrant<br>(Non-Occluded) | 18 | 18 | 18 | 18 | 18 | 18 |
| Lower-left quadrant<br>(Non-Occluded) | 18 | 18 | 18 | 18 | 18 | 17 |
| Lower-right quadrant<br>(Occluded) | 15 | 17 | 9 | 13 | 15 | 7 |

**Table S2.** Subjects with significant individual subject cross-classification (Wilcoxon signed-rank).

|  | V1 |  | V2 |  |
| --- | --- | --- | --- | --- |
|  | Category | Depth | Category | Depth |
| Upper-right quadrant<br>(Non-Occluded) | 18 | 15 | 18 | 15 |
| Lower-left quadrant<br>(Non-Occluded) | 18 | 5 | 18 | 5 |
| Lower-right quadrant<br>(Occluded) | 15 | 5 | 14 | 6 |

